# Bioinformatics analysis of immune-related prognostic genes and immunotherapy in renal clear cell carcinoma

**DOI:** 10.1101/2022.07.22.501084

**Authors:** Ziwen Pan, Sheng Chang, Song Chen, Daqiang Zhao, Zhiyu Zou, Linrui Dai, Yibo Hou, Qianqian Zhang, Yuanyuan Yang, Zhishui Chen, Weijie Zhang, Yuanyuan Zhao

## Abstract

Clear cell renal cell carcinoma (ccRCC) is an immunogenic tumor and investigating the immunorelated genes is of great significance. To investigate the immunoprognostic genes of ccRCC, we analyzed the data assimilated from a public database (The Cancer Genome Atlas (TCGA) database and the gene expression omnibus (GEO) database) using bioinformatics. Then, a immunoprognosis model was constructed to identify 4 hub genes with moderate predictive values for the prognosis of ccRCC patients. Interestingly, these 4 genes were associated with the prognosis of ccRCC patients based on Oncomine and Gena Expression Profiling Interactive Analysis (GEPIA) databases. The correlation analysis between the immune infiltrate, immune checkpoints, and immunotherapy and this immunoprognosis model showed that immune infiltrate and could predict the immunotherapy effects. We also conducted a quantitative real-time polymerase chain reaction analysis and found that the expressions of 3 hub genes were associated with tumor progression (*P*<0.1). In conclusion, 4 genes that may serve as potential biomarkers in ccRCC were identified with respect to prognosis.

## Introduction

ccRCC is an immunogenic tumor and benefits from immunotherapy. Ipilimumab, an anti-CTLA-4 antibody, combined with nivolumab, showed a response rate of about 40% in the CheckMate 016 trial [1]. Recent advances in bioinformatics have focused attention on the correlation between ccRCC prognosis and immunity. The targeted immune cell checkpoints have been proven to be one of the most effective methods to activate an anti-tumor immune response. Cytotoxic T lymphocyte-associated protein 4 (CTLA-4) and programmed cell death protein 1 (PD-1) are two common immune checkpoints. Several studies reported immuoprognostic models (IPMs). Luo et al. [2] revealed three IPMs to assess the prognosis of metastatic colorectal cancer, obtained at 1, 3, and 5 years; the area under the curve (AUC) was 0.646, 0.7, and 0.692, respectively. Moreover, the tumor-infiltrating immune cells (TIICs) [3] are crucial in modulating tumor progression and have a potential prognostic value. A previous study showed that the intratumoral immune cell infiltration is associated with the outcomes of cancer patients. Choueiri et al. [4] reported that kidney cancer patients with high abundance of CD8+T cells have poor prognosis, while Bromwich et al. [5] indicated that CD4+T cell infiltration was associated with poor survival. Presently, the mechanism of how immunity affects the occurrence and development of ccRCC is yet unclear. Thus, the current study aimed to establish a correlation between immunity and the occurrence and development of ccRCC. In this study, we analyzed the immune-related differentially expressed genes (DEGs) associated with ccRCC prognosis. These findings would be beneficial in improving the prognosis of patients with ccRCC.

## Materials and Methods

### Data information and database analysis

Data were obtained from TCGA and GSE53757 of the GEO database [6]. The GSE53757 source files were transformed into log2 and normalized to expression matrices. DEGs were calculated using the R package “Limma” to differentiate between ccRCC and normal kidney samples. Subsequently, the DEGs were screened if the log2fold-change was >1 and the false discovery rate (FDR) was 0.05. TCGA data after pretreatment using the “edgeR” R package for variance analysis were set to several times the expression of genes, the amount of correction after each gene expression to >1, the domain value as |log2fold-change|>2, and FDR<0.05 for screening the DEGs. The “VennDiagram” package was used to identify the intersection of the DEGs obtained from TCGA and GEO database and the immune-related genes (IRGs) downloaded from the Import database.

### Function enrichment analysis of DEGs

We used the R packages of “clusterprofiler,” “enrichplot,” and “ggplot2” for Gene Ontology (GO) and Kyoto Encyclopedia of Genes and Genomes (KEGG) pathway analysis. Significant *P*- and *Q*-values were set to 0.05.

### Prognostic-related survival model

Using IRGS, univariate Cox risk regression and least absolute shrinkage and selection operator (LASSO) regression analysis (Simple Random sampling of “caret” package with R software was performed at the ratio of 2:1 for the training and test groups), multivariate Cox risk regression was analyzed to build a risk prognosis model, and survival-related hub genes were screened out. *P*<0.001 was statistically significant in the univariate Cox regression. The “Glmnet” package of R was used to set 1000-fold replacement and 10-fold cross validation for LASSO regression. Kaplan–Meier survival plots showed the prognosis of ccRCC patients. the AUC was calculated using the R package “survivalROC.”

### Oncomine and GEPIA databases [7] were used to analyze the prognostic value of DEGs

In order to verify the hub genes, five qualified databases were obtained from Oncomine [8] (www.oncomine.org/), followed by meta-analysis. The filter conditions were set as follows: the cancer type was ccRCC, the sample type was mRNA, and the analysis type was case-control study. The threshold was set at 1e-4, the fold-change to 2, and the gene ranked up to 10%. Then, the GEPIA database (http://gepia.cancer-pku.cn/) was analyzed to determine the survival prognosis of hub genes.

### Immunoinfiltration analysis of prognostic genes

Cibersort [9] is a deconvolution algorithm that quantifies the cell components from a large number of tissue gene expression profiles. Because of the prior knowledge of the expression profiles of purified leukocyte subpopulations, Cibersort can estimate the immune composition of the tumor tissue accurately.

### Genetic alteration in genes key related to prognosis

The mutation data were obtained from TCGA database. Next, we analyzed the genetic changes in different risk groups and their correlation with prognosis.

### Immunotherapy and immune targets of prognostic genes associated with ccRCC

We investigated the differential expression of *PD-1* and *CTLA-4* genes in patients in different risk groups. In addition, we downloaded 4 (kidney renal clear cell carcinoma) KIRC files related to immunotherapy from the cancer immunoAtlas [10] (https://tica.at/). These included ctla4_neg_pd1_neg, ctla4_neg_pd1_pos, ctla4_pos_pd1_neg, and ctla4_pos_pd1_pos. Under the four conditions above, the Wilcoxon test was used to evaluate the efficacy of immunotherapy in the two risk groups.

### RNA extraction and quantitative real-time polymerase chain reaction (qRT-PCR)

A total of 12 pairs of ccRCC and para-cancerous tissues were collected from Tongji Hospital of Huazhong University of Science and Technology (Wuhan, China) between January 2022 and May 2022. Total RNA was isolated from ccRCC and normal kidney cells using RNA-easyTM Isolation Reagent Vazyme Cat (Vazyme, Nanjing, China). A reverse transcription kit (Vazyme) was used to convert the mRNA into cDNA. qRT-PCR was carried out with a SYBR Green kit R701-02 (TransGen Biotech, Beijing, China) in a final volume of 10 mL to detect the expression level of these genes. The data were analyzed using StepOne software (Thermo Fisher Scientific). *GAPDH* was used as a reference gene to calculate the relative expression levels using 2-DDCT. Primer sequences are listed in S1 Table.

### Statistical analysis

R package “edgeR” and “Limma” were used to analyze the differential genes. Two groups were compared using Wilcoxon test. Survival analysis was carried out using Kaplan–Meier method and log-rank test. The survival-related receiver operating characteristic (ROC) package was utilized to draw the ROC curve. Chi-square test was used to analyze the differences between two groups in terms of clinical parameters, and the difference was considered statistically significant at *P*<0.05. All statistical analyses were performed using GraphPad Prism 9 software (GraphPad Software, San Diego, CA, USA).

## Results

### Characteristics and functional clustering of 128 differentially expressed IRGs in ccRCC

A total of 1777 DEGs were obtained from the TCGA database (Fig 2a), and 2938 DEGs were obtained from GEO database (Fig 2b). These were pooled with 2483 IRGs, and finally 128 IRGs (Fig 2c and Table 1) overlapping in both databases were identified, including 102 upregulated and 26 downregulated IRGs. GO (Fig 2d) and KEGG (Fig 2e) enrichment analyses of the above IRGs showed a response to chemokines, external side plasma membrane, and receptor-ligand activity, which are the most common biological terms for BP, CC, and MF. The most enriched form in KEGG pathway analysis was the cytokine-cytokine receptor interaction. The study schematic is illstrated in Fig 1.

**Table 1:**
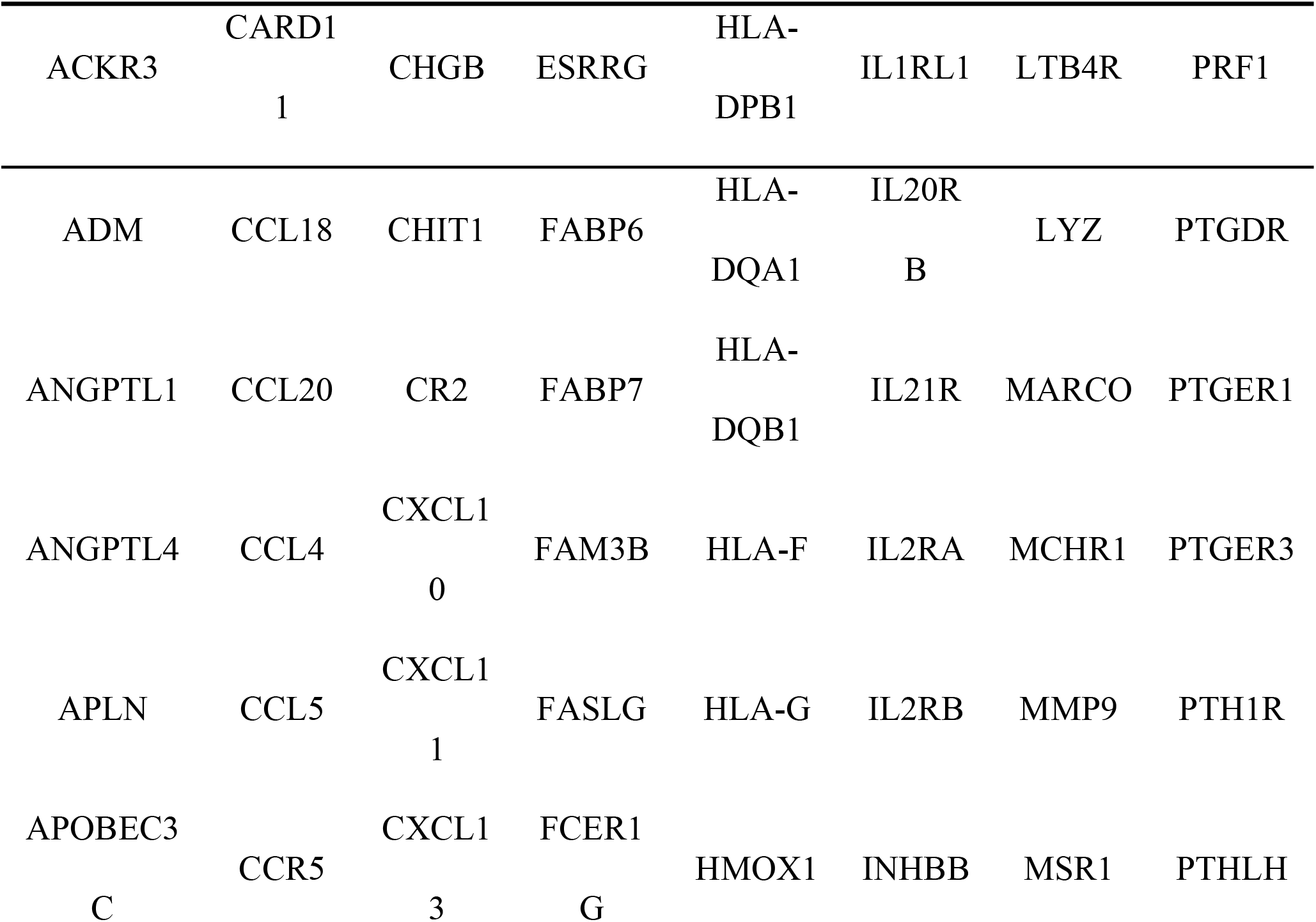
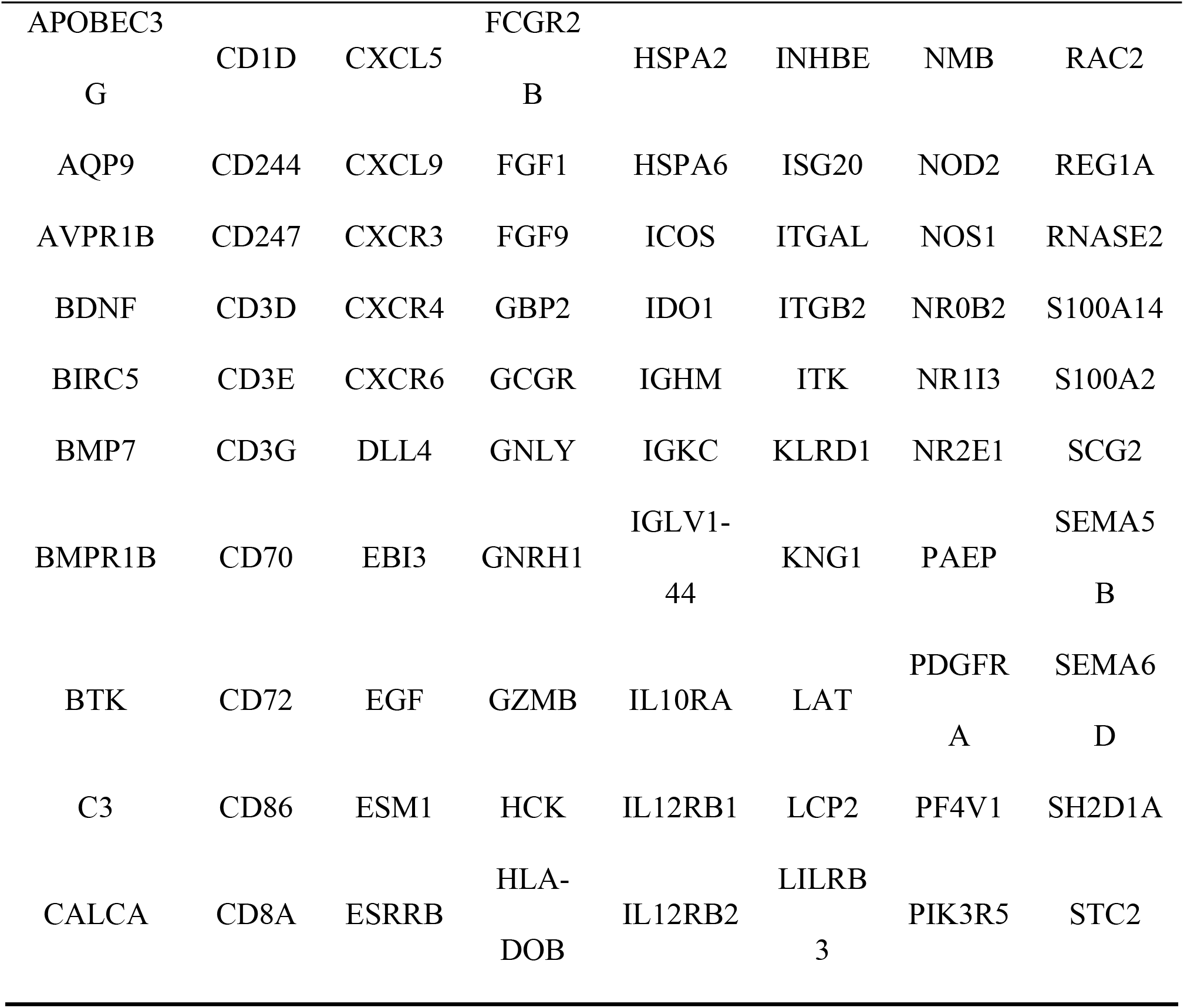
128 IRGs.

**Fig 1:**
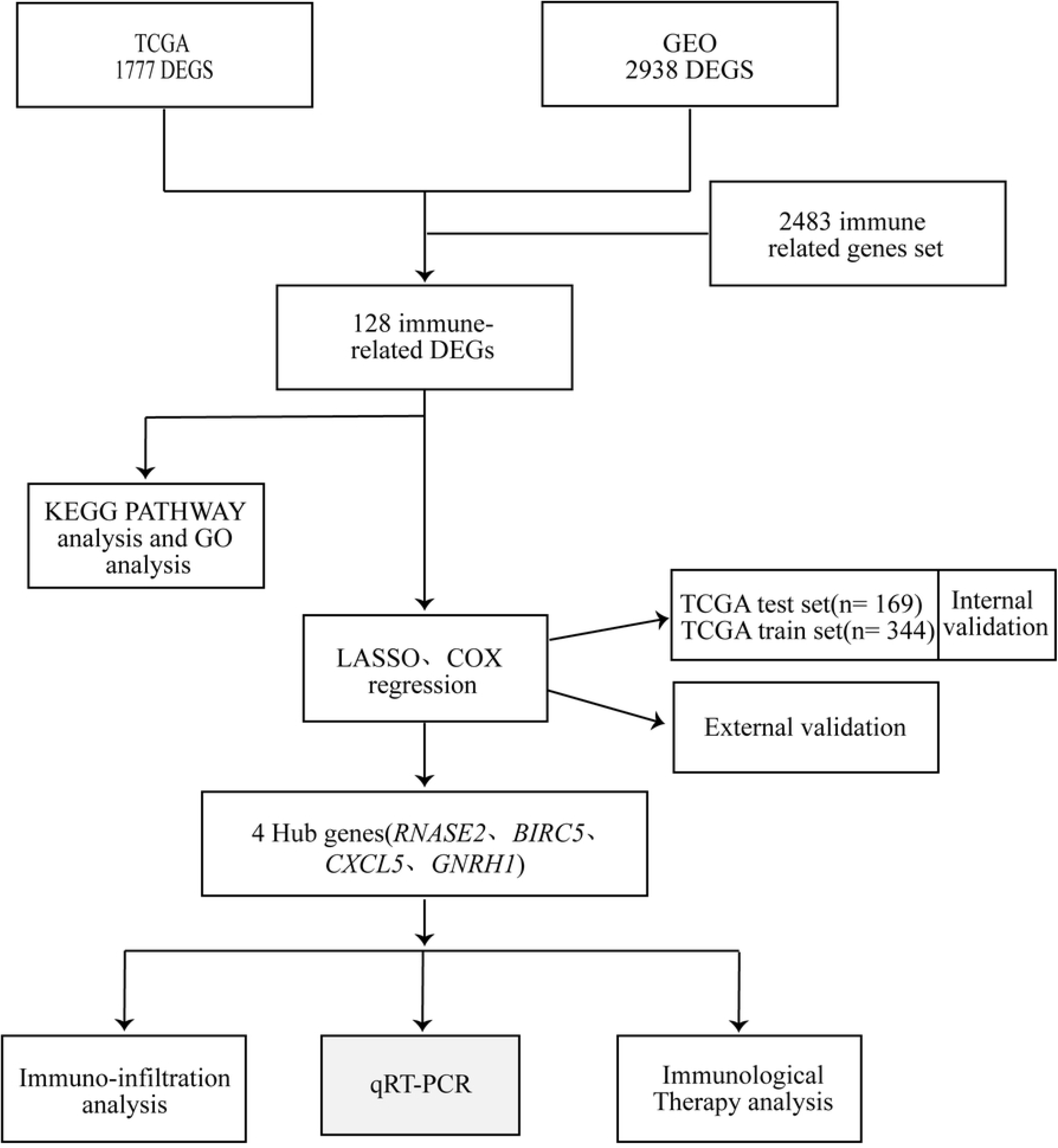
Schematic of the study process.

**Fig 2:**
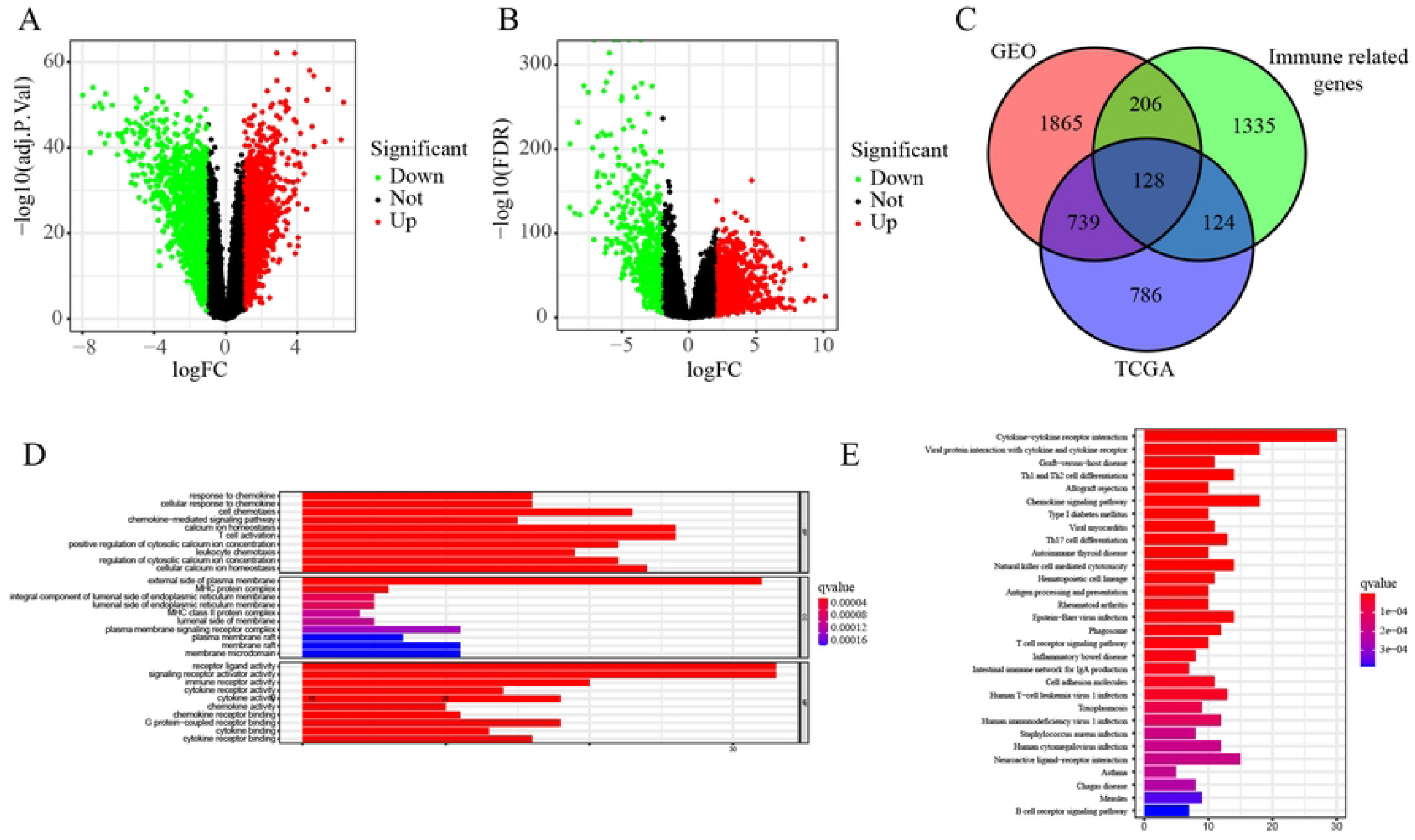
Differentially expressed IRGs. (a-b) represents the volcano map of DEGs from the TCGA and GSE53757 datasets. (c) Venn diagrams of DEGs in TCGA and GSE537357 dataset and IRGs. (d-e) GO and KEGG enrichment analysis of differential expression of IRGs. BP, biological process. CC, cellular component; MF, molecular function.

**Figs 1:** High- and low-risk grouping and clinical traits. (a–d) Boxplot representing risk scores for different clinical traits.

### Establishment and validation of prognostic risk models of the 4 hub genes

In order to identify the key IRGs in the occurrence and development of ccRCC and use them as potential predictive targets, 128 IRGs were evaluated by univariate Cox regression analysis. Among these, 40 DEGs related to survival were screened out, and a forest plot was constructed (*P*<0.001; Fig 3c). LASSO regression analysis (Fig 3a and 3b) was performed, and 5 DEGs were obtained. Then, multivariate Cox regression analysis was performed, and 4 prognostic genes were screened out: *BIRC5, GNRH1, RNASE2*, and *CXCL5* (Table 2). Immune-related gene-based prognostic index (IRGPI) was calculated according to the risk score (Table 2). The risk score of patients was determined by the following formula:

**Table 2:** Multivariate Cox regression analysis revealed 4 prognostic markers genes.

| Gene symbol | Coef | HR | HR.95L | HR.95H | Pvalue |
| --- | --- | --- | --- | --- | --- |
| BIRC5 | 0.219681 | 1.24568 | 1.035347 | 1.498741 | 0.019907 |
| GNRH1 | 0.492465 | 1.636344 | 1.301839 | 2.056801 | 2.44E-05 |
| RNASE2 | 0.383803 | 1.467856 | 1.147024 | 1.878426 | 0.002288 |
| CXCL5 | 0.096701 | 1.101531 | 1.007325 | 1.204547 | 0.034009 |

**Fig 3:**
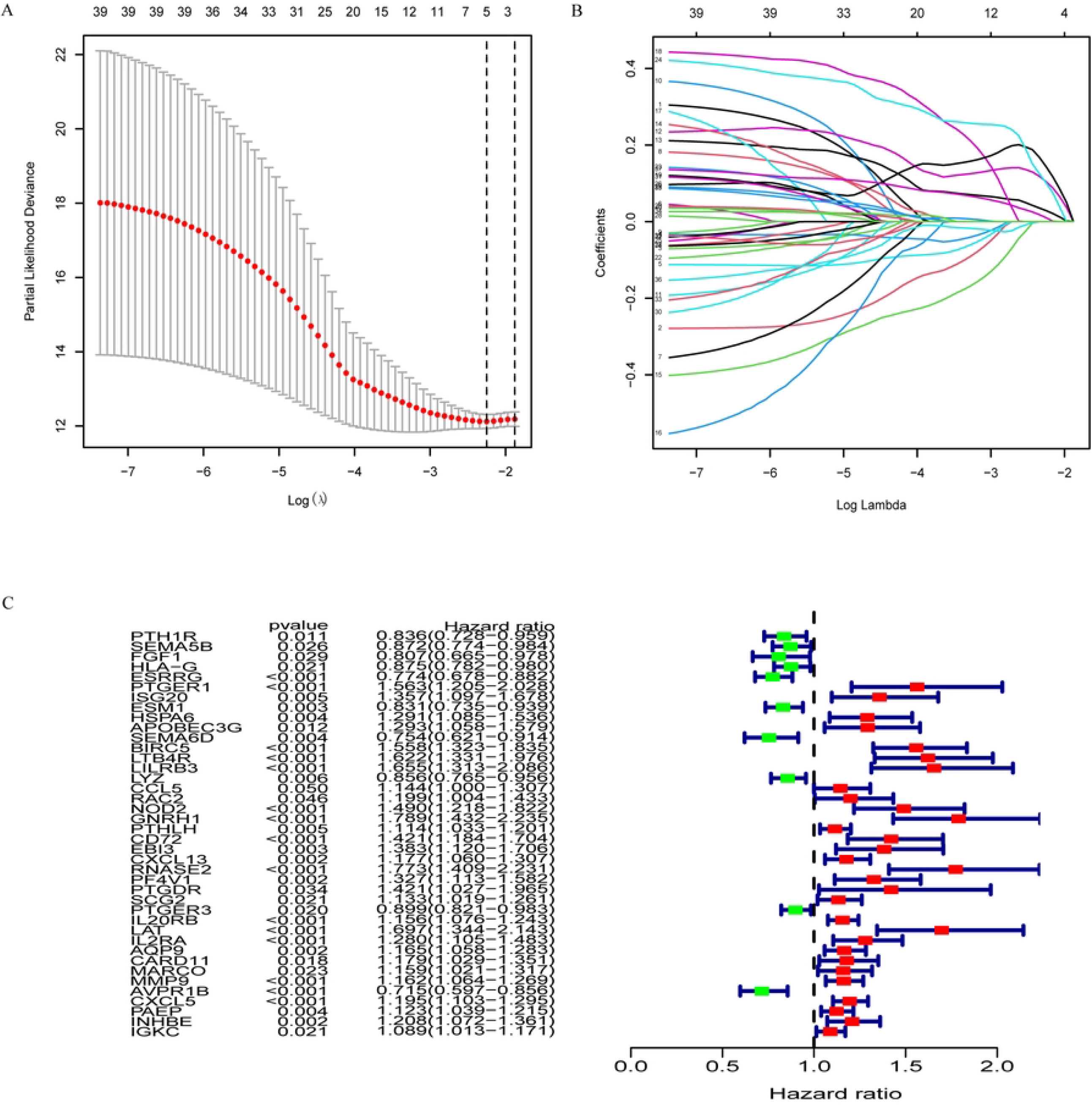
LASSO regression and univariate Cox regression. (a-b) LASSO regression model. (c) Forest plot consists of 21 IRGs.

Risk

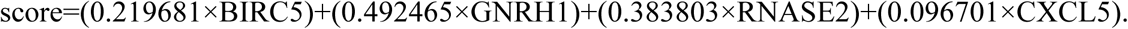

The median of the risk score calculated by IRGPI-based risk prognosis model was 0.9243977, and the samples downloaded from TCGA database were divided into the low- and high-risk groups. To verify the reliability of the model, we analyzed the outcomes and found that the low-risk group had better outcomes than the high-risk group (*P*<0.001; Fig 4a–4c). The ROC curve showed that our models have a moderate predictive value. We also analyzed the correlation between 4 hub genes and clinically relevant indicators and found that the gender, tumor stage, and tumor grade were correlated with risk score (*P*<0.05; Figs 1b–1d). Additionally, the 4 hub genes were significantly upregulated in the tumor group based on the GEO database (*P*<0.001; Fig 5e). Then, we conducted a meta-analysis of 4 hub genes based on the results from the Oncomine database, and the results were consistent with previous results (*P*<0.05; Fig 5a–5d). To verify whether these 4 hub genes are related to prognosis, survival analysis was performed using GEPIA tools (http://gepia.cancer-pku.cn/); the expression of the genes was negatively correlated to tumor prognosis (*P*<0.05; Fig 5f–5i).

**Fig 4:**
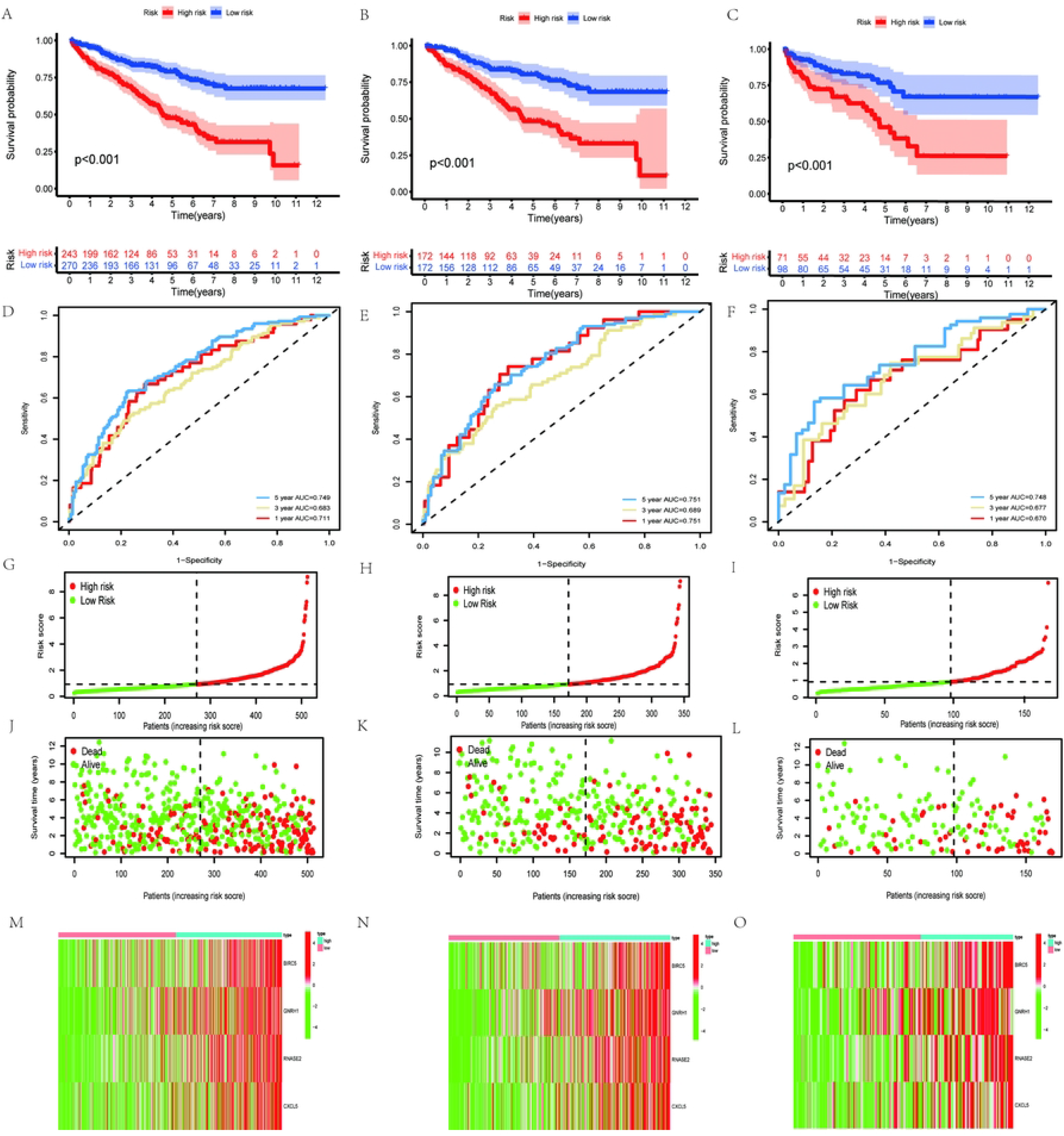
Survival models and internal validation of prognostic IRGPI for all patients with ccRCC, training and test groups, and correlation between risk scores and clinical traits. (a–c) Three groups of patients had good prognosis in the low-risk group, respectively. (d–f) ROC curves of survival prognostic model of the three groups. (g–i) risk score of the high and low-risk group in the three groups. (j–l) survival time and status of patients in the three groups. (m–o) was the heat map of the 4 hub genes between the high- and low-risk groups.

**Fig 5:**
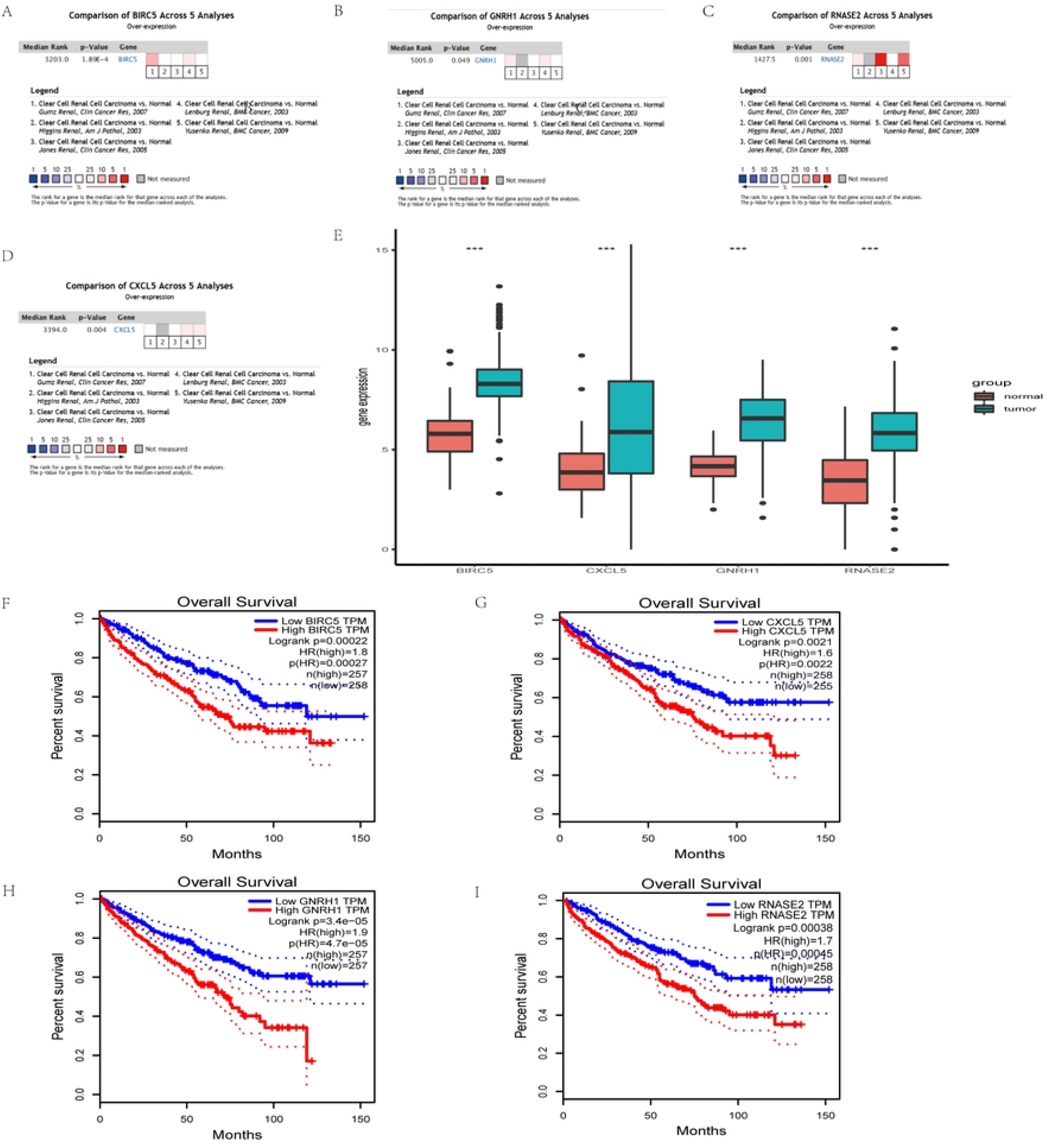
Differential expression and external validation of 4 hub genes between tumor and normal groups. (a–d) external validation using the Oncomine database. (e) boxplot of the differential expression of 4 hub genes in TCGA. (f–i) expression of the genes was negatively correlated to tumor prognosis (\*\*\**P*<0.001).

### Characteristics of immune invasion in the two risk groups were significantly different

We analyzed the prognosis of immune cell content in the two risk groups (Fig 6a–6d) and found that patients with high resting mast cells, low plasma cells, low follicular helper T cells, and low regulatory T cells showed improved prognosis.Moveover,The evaluation of the two risk groups of patients and correlation immunity infiltration (Fig 6e) revealed that high-risk group had higher levels of CD4 memory T cells, follicular helper T cells, and regulatory T cells, while the low-risk group had lower levels of resting dendritic cells, resting mast cells, and neutrophils. Moreover,. These data suggested that the immune status of ccRCC can be predicted based on 4 hub genes.

**Fig 6:**
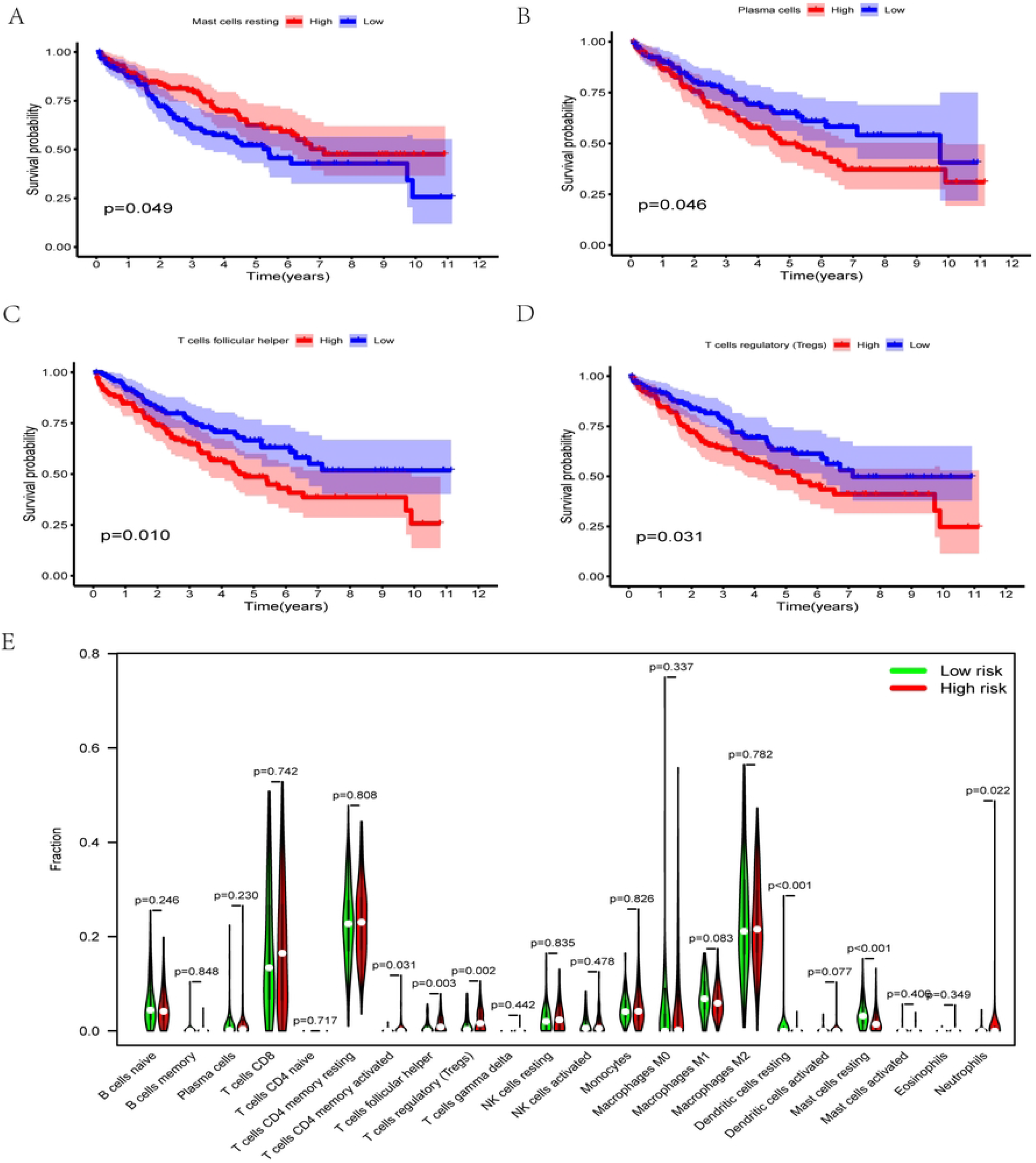
A comparison between two risk groups and the immune cell content and prognosis of immune cells. (a–d) Prognostic analysis of patients with a varying number of resting mast cells, plasma cells, follicular helper T cells, and regulatory T cells. (e) difference analysis of immune cell content in the two risk groups.

**Figs 2:** High- and low-risk assessment grouping and genetics changes. (a-b) Waterfall plot of tumor mutation load in the two risk subgroups. (c) represents the difference analysis of tumor mutation load in the two risk subgroups.

### Mutation load was significantly different between the two risk groups

A high mutation load and increase in neoantigens were associated with the clinical effects of the immunocheckpoint inhibitor therapy. Therefore, we analyzed the genetic changes in the two risk groups of samples (Figs 2a and 2b) and found the primary mutation in the high-risk group in Missense_Mutation, In_Frame_Del, Frame_Shift_Ins, Translation_Start_Site, Frame_Shift_Del, In_Frame_Ins, Nonsense_Mutation, and Multi_Hit in three genes, *VHL, PBRM1*, and *SETD2*. The main mutation points in the low-risk group were Nonsense_Mutation, Translation_Start_Site, Frame_shift_Del, In_Frame_DeI, Frame_Shift_Ins, Nonstop_Mutation, Missense_Mutation, and Multi_Hit in three genes: *VHL, PBRM1*, and *TTN*. We also analyzed the differences in the tumor mutation load expression in the two groups (Figs 2c) and found that the high-risk group had a higher tumor mutation load than the low-risk group.

### Expression of immunotarget-related genes and the efficacy of immunotherapy differ significantly in the two risk groups

We analyzed the differential expression of *PD-1* and *CTLA-4* in both groups and found that *CTLA-4* was overexpressed in the high-risk group, while *PD-1* was overexpressed in the low-risk group (Fig 7a and 7b). Next, to demonstrate the effectiveness of our survival prognosis model, we analyzed the efficacy of immunotherapy in the two risk groups under four different conditions (Fig 7c–7f). The immunotherapy efficacy of patients in the high-risk group was significantly stronger when they received either CTLA-4-positive or both treatments. The findings indicated that immunotherapy is more effective in high-risk ccRCC than in the low-risk group.

**Fig 7:**
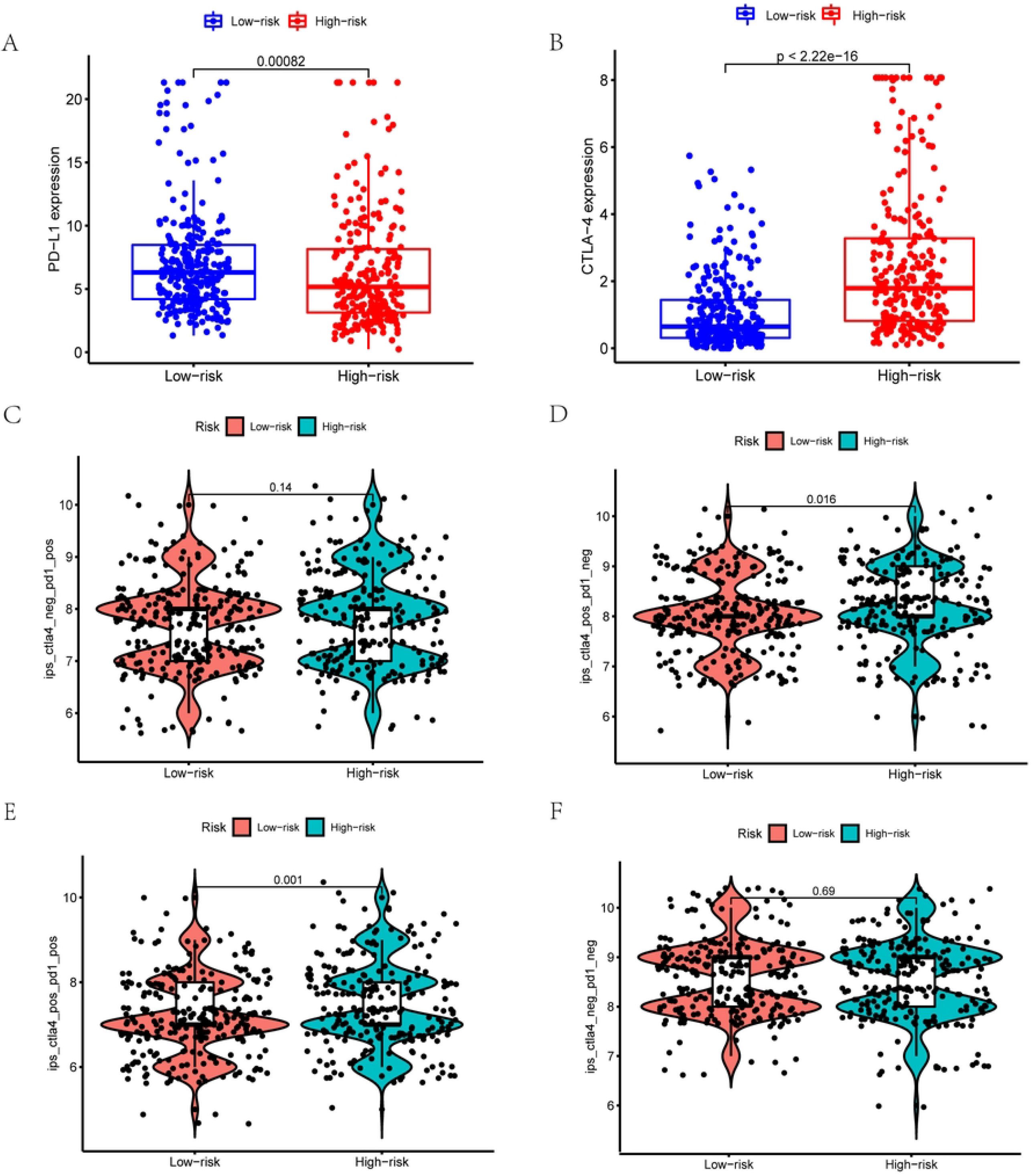
Differential expression of *PD-1* and *CTLA-4* and related immunotherapy in patients in the two risk groups. (a-b) represents the differential expression of *PD-1* and *CTLA-4* in the two risk groups. (c–f) represents the difference analysis of patients in the two risk groups when receiving immunotherapy for anti-PD-1 and CTLA-4.

### Expression of 4 hub genes by qRT-PCR at the cellular and tissue levels of ccRCC

According to qRT-PCR analysis, *BIRC5* and *CXCL5* mRNA levels were significantly higher in ccRCC than paracancer tissues (*P*<0.05; Fig 8b-8c). Furthermore, compared to the normal renal epithelial cell line HK-2, the ccRCC cell line 786O expressed significantly more *BIRC5* (*P*<0.01; Fig 8f). Due to the short postoperative follow-up time of the collected cases, we analyzed the correlation between the qRT-PCR expression values of the 4 hub genes and clinicopathology (Table 3) and found that the expression values of 3 hub genes (*RNASE2, BIRC5*, and *GNRH1*) were associated with tumor progression (*P*<0.1).

**Table 3:** The correlation between the expression levels of 4 hub genes and clinicopathology.

| Characteristic | RNASE2 |  | P<br>value | BIRC5 |  | P<br>value | CXCL5 |  | P<br>value | GNRH1 |  | P<br>value |
| --- | --- | --- | --- | --- | --- | --- | --- | --- | --- | --- | --- | --- |
|  | expression |  |  | expression |  |  | expression |  |  | expression |  |  |
|  | high | low |  | high | low |  | high | low |  | high | low |  |
| ISUP grade |  |  |  |  |  |  |  |  |  |  |  |  |
| 1 | 2 | 4 | >0.1 | 1 | 5 | >0.1 | 2 | 4 | >0.1 | 2 | 4 | >0.1 |
| 2-4 | 3 | 3 |  | 2 | 4 |  | 2 | 4 |  | 3 | 3 |  |
| pT |  |  |  |  |  |  |  |  |  |  |  |  |
| T1 | 5 | 4 | <0.1 | 3 | 6 | <0.1 | 3 | 6 | >0.1 | 5 | 4 | <0.1 |
| T2-T3 | 0 | 3 |  | 0 | 3 |  | 1 | 2 |  | 0 | 3 |  |
| PN |  |  |  |  |  |  |  |  |  |  |  |  |
| N0 | 4 | 6 | >0.1 | 2 | 8 | >0.1 | 4 | 6 | >0.1 | 5 | 5 | >0.1 |
| N1 | 1 | 1 |  | 1 | 1 |  | 0 | 2 |  | 0 | 2 |  |
| Pm |  |  |  |  |  |  |  |  |  |  |  |  |
| M0 | 5 | 6 | >0.1 | 3 | 8 | >0.1 | 4 | 7 | >0.1 | 5 | 6 | >0.1 |
| M1 | 0 | 1 |  | 0 | 1 |  | 0 | 1 |  | 0 | 1 |  |

ISUP : International Society of Urological Pathology

**Fig. 8:**
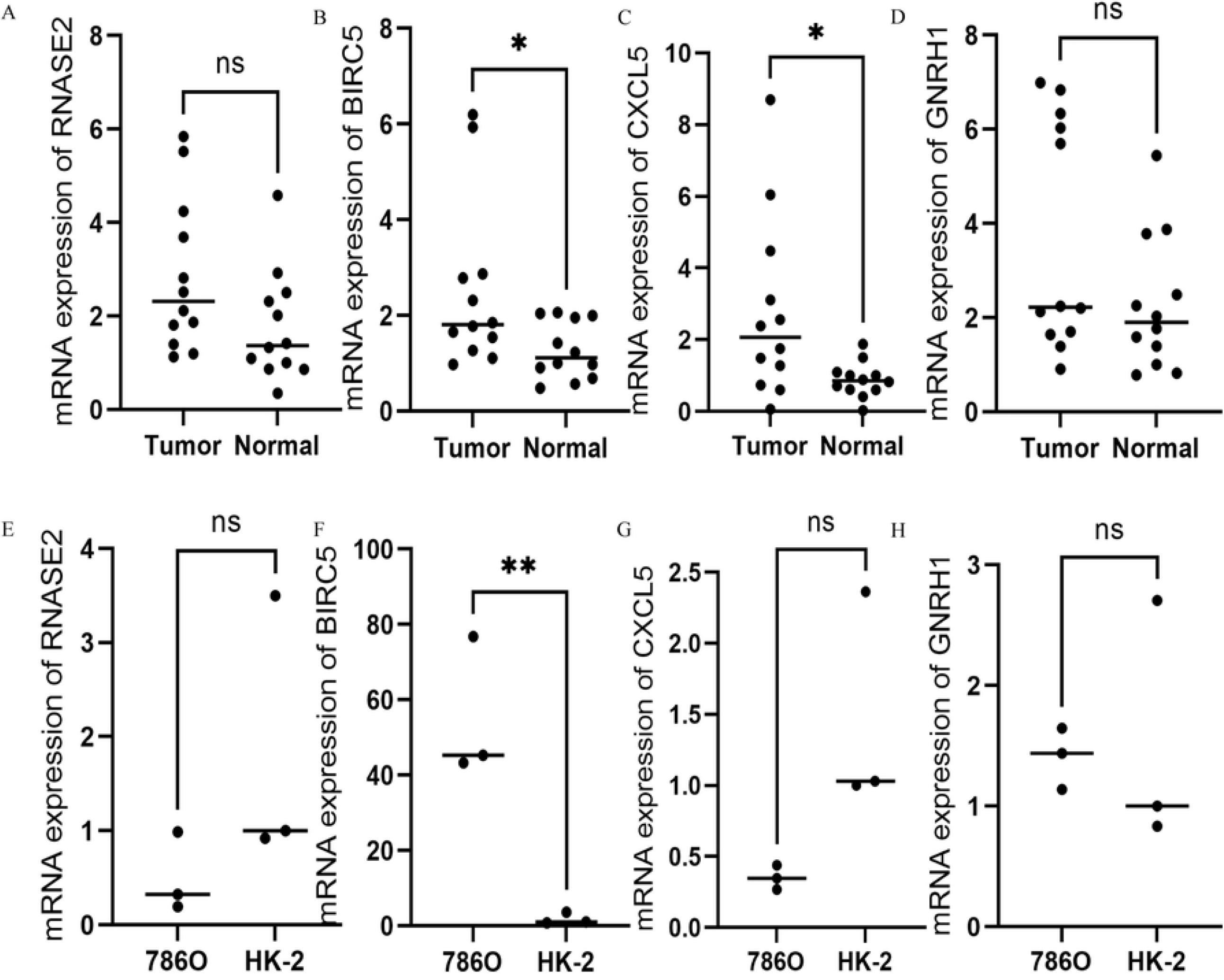
The mRNA levels of the 4 hub genes. (a–d) Expression of the 4 genes in 12 pairs of ccRCC and para-cancer tissues by qRT-PCR. (e–h) expression of the 4 hub genes in cancer cells and normal kidney cells by qRT-PCR (\*\**P*<0.01; \**P*<0.05).

## Discussion

A total of 128 IRGs differentially expressed in ccRCC tissues compared to normal tissues were identified from TCGA and GEO databases. Subsequently, functional enrichment analysis found that these genes were mainly involved in inflammation and immunity. Univariate Cox, LASSO, and multivariate Cox regression analyses revealed 4 prognostic hub genes (*RNASE2, BIRC5, CXCL5*, and *GNRH1*), and a prognosis model was established. We also verified the reliability of the model and found that the 4 hub genes were closely related to the prognosis of ccRCC patients. To further analyze the reliability of the model, we evaluated the expression of immune infiltrate and tumor mutation load, IRGs, and immunotherapy effect in the two risk groups and found that the 4 hub genes could predict the immune status of ccRCC patients. These results demonstrated that *BIRC5* expression was significantly lower in ccRCC and tumor cell lines than in paracancer tissues and normal cell lines. The expression of *CXCL5* was significantly higher in tumor tissue than paracancer tissues; however, the expression of *CXCL5* in 786O and HK-2 cells did not differ significantly, which might be because the cancer cell lines do not represent the gene expression of tumor tissues. Also, *GNRH1* and *RENSE2* expressions did not differ significantly between the tissues and cells. Next, we evaluated whether the expression values of these hub genes could affect the prognosis. Due to the short follow-up period, we analyzed the correlation between clinicopathology and the gene expression values and found that tumor grade was correlated with the expression values of the 3 hub genes. However, due to the small sample size and short follow-up time, the association was not statistically significant (*P*>0.05), necessitating further exploration.

*BIRC5* (Survivin), is strongly expressed in most human tumors and closely related to tumor progression, recurrence, and chemotherapy resistance [11]. It is also associated with poor prognosis and progression of ccRCC [12,13]. Current studies have shown that *RNASE2*, a member of the pancreatic ribonuclease family, is a useful prognostic predictor of ccRCC [14]. Among cancer patients, *CXCL5* [15] is closely related to the survival time, recurrence, and metastasis and is associated with a poor prognosis in ccRCC [16]. Microslaw et al. [17] demonstrated that the expression levels of chorionic gonadotropin beta subunit and *GnRH1* in the blood of cancer patients might be valuable in indicating tumor metastasis and spread. Wu et al. [18] demonstrated that *GnRH1* and *LTB4R* are prognostic biomarkers for patients with ccRCC.

Reportedly, a risk model can predict the prognosis of patients. For example, Hua et al. [19] combined multiple databases with building a survival model of immune-related prognostic genes, which could accurately predict the prognosis of ccRCC patients. Wan et al. [14] used 7 IRGs to establish a regression model for predicting the prognosis of ccRCC patients and found that the risk model could accurately distinguish patients with different survival outcomes and identify those with high and low mutation loads. Notably, our model could also predict patient prognosis. The 4 hub genes were regarded as independent prognostic factors related to the clinicopathological characteristics.

In addition, we analyzed the correlation between the risk model and immune filtration, immune target, and immunotherapy and found that the 4 genes predicted the immune status. These analyses demonstrated a poor response of the high-risk group to immunosuppression.

In a study by Zhang et al. [20] low levels of resting dendritic cell infiltration were observed in the high mutation load (TMB) group, and high levels of TMB were associated with low survival rates while low resting dendritic cell infiltration was associated with poor prognosis. Castle et al. [21] reported that dendritic cells activate T cells in patients and increase cytotoxic T cell production, exerting anti-tumor effects. Basile et al. [22] demonstrated that high tumor-infiltrating neutrophils were linked to poor overall survival in ccRCC patients. A previous study found that in patients with ccRCC mast cells [23] and high infiltration, the immune treatment may be effective with an improved prognosis. Plasma cells and T cells are crucial in tumor immunity. In the current study, neutrophils, CD4 memory T cells, follicular helper T cells, and regulatory T cells were abundant in the high-risk group, while resting mast cells were low. We further analyzed the prognosis of patients with varying content of immune cells and found that the patients with a high number of resting mast cells had better outcomes. On the other hand, the patients with low plasma cells, follicular helper T cells, and regulatory T cells had a poor prognosis.

Previous studies showed that the tumor mutation load is positively related to the presence of a novel antigen that significantly enhances the efficacy of immunotherapy. Subsequently, we tested our model and found that the high-risk group had a high mutation load; these results were in agreement with those of Hua et al. [19]. According to the immunotherapy results, we found that the high-risk group had better immunotherapy efficacy. These observations further confirmed the functional correctness of our prognosis model based on the 4 hub genes.

However, the current study differs from the previous studies in some aspects. First, we combined multiple databases to construct the survival models of 4 hub genes related to prognosis and performed the internal and external validation, and qRT-PCR results showed that the two hub genes were differentially expressed in the tumor and normal tissues. Subsequently, we analyzed the immune cell content in different risk groups and the prognosis of patients with varied immune cell content. Finally, we analyzed the differential expression of immune targets, *PD-1* and *CTLA-4*, in the two risk groups in the survival model constructed using the 4 hub genes and verified the data related to the treatment using the two immune targets downloaded from the public database, proving the accuracy of the model function.

Nevertheless, this study has some limitations. The first is the limited sample size; hence, the findings requires further substantiation using multiple samples and multicenter clinical trials. Second, the exact mechanism by which these 4 genes influence the prognosis of ccRCC patients is yet unknown. In future studies, we need to follow-up ccRCC patients prospectively and detect the correlation of these 4 genes to confirm the accuracy of the predictive model on the occurrence, development, and prognosis of ccRCC.

## Conclusions

We used 4 hub genes to construct a prognostic risk model to verify its accuracy. As a prognostic marker, the risk score of this model might distinguish the immune status, immunotherapy effect, and prognosis of patients with different risk scores.

## Acknowledgements

We would like to thank the authors who generously shared their data and all study participants and the anonymous reviewers for their useful comments on the manuscript.

## Supplementary Material

**S1Table. Primer sequences**.

**S1 Fig. High- and low-risk grouping and clinical traits**.

**S2 Fig. High- and low-risk assessment grouping and genetics changes.**

**Ethical approval**

